# Semi-automated workflow for high-throughput *Agrobacterium*-mediated plant transformation

**DOI:** 10.1101/2024.10.09.617252

**Authors:** Davide Annese, Facundo Romani, Carolina Grandellis, Lesley Ives, Eftychios Frangedakis, Felipe X. Buson, Jennifer C. Molloy, Jim Haseloff

## Abstract

High-throughput experiments in plants are hindered by long generation times and high costs. To address these challenges, we present an optimized pipeline for *Agrobacterium tumefaciens* transformation and simplified a protocol to obtain stable transgenic lines of the model liverwort *Marchantia polymorpha*, paving the way for efficient high-throughput experiments for plant synthetic biology and other applications. Our protocol involves freeze-thaw *Agrobacterium* transformation method in 6-well plates that can be adapted to robotic automation. Using the Opentrons open-source platform, we implemented a semi-automated protocol showing similar efficiency compared to manual manipulation. Additionally, we have streamlined and simplified the process of stable transformation and selection of *M. polymorpha*, reducing cost, time, and manual labour without compromising transformation efficiency. The addition of sucrose in the selection media significantly enhances the production of gemmae, accelerating the generation of isogenic plants. We believe these protocols have the potential to facilitate high-throughput screenings in diverse plant species and represent a significant step towards the full automation of plant transformation pipelines. This approach allows testing ∼100 constructs per month, using conventional plant tissue culture facilities. We recently demonstrated the successful implementation of this protocol for screening hundreds of fluorescent reporters in *Marchantia* gemmae.

## INTRODUCTION

Plant biotechnology and synthetic biology have rapidly advanced, aiming to genetically enhance crops, improve disease and pest resistance, and boost nutritional value and yields (Yang and Reyna-Llorens, 2023). To achieve these goals, hundreds of genetic parts need to be tested in a high-throughput and cost-effective manner to build comprehensive atlases or toolsets. However, working with plants for genetic engineering presents significant challenges compared to model organisms like bacteria or yeast. While heterologous or cell-free systems can be convenient for high-throughput experimentation, many biological approaches require working *in planta*. Current high-throughput plant transformation strategies involve infiltration of vegetative tissues with *Agrobacterium tumefaciens* (*Agrobacterium* for short), particle bombardment, or osmotic-based methods in protoplasts. Among them, agroinfiltration of *Nicotiana benthamiana* leaves is popular, but transient systems are also applicable to leaf discs and other model plants (Zhang et al., 2020). However, transient transformation has limitations for studying genetics in a developmental or physiological context and scalability needs expensive special equipment (Reed et al., 2017). On the other hand, generating stable and isogenic transgenic plants is time-consuming due to long generation times and the need for expensive plant growth facilities (Ayub and Soto, 2023).

To speed-up design-build-test-learn cycles for plant biotechnology, automation and optimization of different steps of plant transformation are necessary. Although *Agrobacterium*-based transformation methods are widely used for plant transformation, protocols for the transformation of the bacteria are often overlooked in optimization efforts and no automated protocols are available. The most commonly used protocol, electroporation, yields high transformation efficiency (Wise et al., 2006) but is challenging to scale up and can be expensive due to the use of 25-well and 96-well electroporation plate systems (Buchser et al., 2006). This limits the feasibility of transforming arrayed libraries for large scale screens. Other alternatives, such as chemical transformation, triparental mating, and freeze-thaw methods, are generally cheaper, but less effective than electroporation (Wise et al., 2006). Among them, the freeze-thaw method is generally simpler, allows transformation in bulk, and more cost-effective than other methods and has potential for scale-up. However, a high-throughput *Agrobacterium*-mediated plant transformation has not been implemented yet.

The liverwort *Marchantia polymorpha* (*Marchantia* for short) has emerged as a valuable model organism for evo-devo studies and synthetic biology (Sauret-Gueto et al., 2020; Bowman et al., 2022). It features a relatively small (280 Mbp) and well characterised genome, a growing community of researchers and well-established genetic engineering tools (Ishizaki et al., 2016; Frangedakis et al., 2021). The haploid nature of the vegetative body of *Marchantia*, extraordinary regenerative ability, and short vegetative life cycle enable the quick generation of stable transgenic lines, compared to obtaining homozygous transgenic lines in flowering plant model systems such as *Arabidopsis thaliana* or *Nicotiana benthamiana*.

*Marchantia* nuclear transformation can be achieved using particle bombardment or *Agrobacterium*-based methods, with the latter being the easiest and most efficient method so far (Chiyoda et al., 2008). Various tissues of *Marchantia*, such as gemmae, adult thalli or spores, can serve as starting materials for plant transformation (Ishizaki et al., 2008; Kubota et al., 2013; Tsuboyama and Kodama, 2018). Sporelings offer an extraordinary tissue for transformation, as a single sporangium can yield 600-1000 independent transgenic plants (Ishizaki et al., 2008). While some efforts were made to simplify *Marchantia* transformation with reasonable efficiency (Tsuboyama and Kodama, 2014; Sauret-Gueto et al., 2020), there has been limited comparison of these protocols.

Considering the need for improved plant transformation methods to enable high-throughput approaches, we optimized and miniaturized a protocol for *A. tumefaciens* transformation based on the classic freeze-thaw method (Hofgen and Willmitzer, 1988). Additionally, several steps of the process were automated using the open-source platform Opentrons OT-2 (https://opentrons.com), allowing up to 96 transformations per batch. The resulting bacterial strains can be then used for either plant transient or stable transformation, depending on the application. Furthermore, we compared different methods for *Marchantia* sporeling transformation and proposed a simplified and miniaturized protocol using 6-well plates (Sauret-Gueto et al., 2020) with optimized selection and rapid generation of propagules. This end-to-end pipeline significantly simplified the process, reducing hands-on work and costs, as well as facilitating scale up without significantly comprising transformation efficiency. The total time required, starting from a genetic construct to obtain a stable transgenic plant ready for downstream analysis, has been reduced to only 4 weeks. We successfully applied this pipeline to test the expression patterns of fluorescent reporters for a collection of over ∼360 promoters, representing approximately 80% of transcription factors in the *Marchantia* genome (Romani et al., 2023). With this approach, it is feasible to test about one hundred constructs per month using conventional plant tissue culture facilities in a cost-effective manner. For both methods, we provide detailed step-by-step protocols and instructional videos to illustrate the process.

## RESULTS

### Transformation of Agrobacterium tumefaciens

To accelerate the transformation of *A. tumefaciens*, we optimized and miniaturized a freeze-thaw transformation protocol (Hofgen and Willmitzer, 1988; Weigel and Glazebrook, 2006). Compared to most existing approaches for the preparation of electrocompetent cells, this method is simpler, cheaper, less laborious and more suitable for large scale experiments. Competent cells were generated by simply growing cells in LB media overnight, concentrating them through centrifugation (see Supplemental Document 1) and finally aliquoting 50 μL in 200 μL tube strips (PCR tubes) or in 96-well plates. These stocks can be stored in a -70/80 °C ultra freezer for several months.

For transformation, 2 μL of miniprep DNA (∼200 ng) of the desired plasmid was added to the ice-thawed competent cells and then flash-frozen in liquid nitrogen for ∼10 seconds. The cells were then transferred to a thermal cycler programmed to deliver a heat shock of 5 minutes at 37°C, followed by 60 minutes at 28 °C for recovery. The transformed cells (50 μL) were then directly plated on 6-well plates containing LB agar with antibiotics for selection (Figure 1A,E). A gentle circular movement was sufficient to spread the cells across the well, eliminating the need for a sterile spatula or glass beads (Supplemental Video 1). The plates were then incubated at 28 °C. Colonies were visible after 2 days, but we recommend growing them for up to 3 days to obtain large colonies.

**Figure 1.**
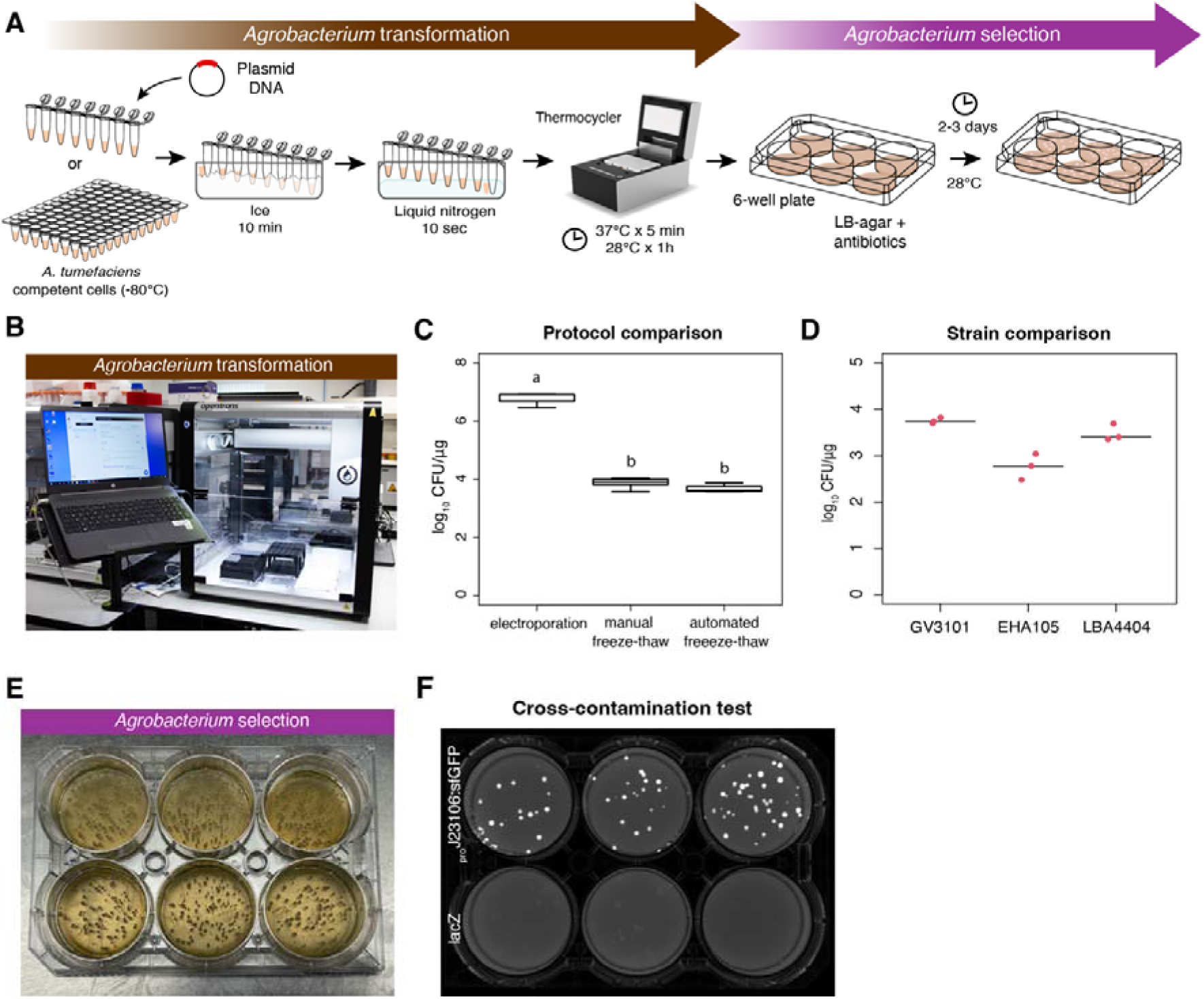
Transformation of Agrobacterium. (A) Workflow schematic diagram of our simplified Agrobacterium freeze-thaw transformation method using PCR tube strips and 6-well plates. (B) The Opentrons OT-2 robot set-up for the automated protocol for Agrobacterium transformation. (C) Comparison of efficiencies for different transformation protocols. (D) Comparison of freeze-thaw transformation efficiency in different Agrobacterium strains. (E) Example of Agrobacterium colonies after selection in a 6-well plate, three days incubation at 28 °C. (F) 6-well plate cross-contamination test showing colonies after selection. Two plasmids expressing a sfGFP or lacZ were alternated and then plated.

Compared to previous applications, our method significantly reduced the bacterial cell volume (from 2-0.25 mL to 50 μL per reaction), DNA amount (from 1-5 μg to 200 ng), and simplified and shortened various steps (see Supplementary Document 1 for details). Instead of using a water bath and a shaking incubator for the heath-shock at 37 °C and subsequent incubation at 28 °C, we found that a thermocycler performed both steps without compromising efficiency. Additionally, the use of 6-well plates proved to be cost-effective, convenient for storage, and amenable to automation of subsequent steps. The transformation efficiency of *Agrobacterium* was determined to be an average of 8 × 10^3^ colony forming units (CFU)/μg DNA (Figure 1C) for a 7.2 kb plasmid, without false positives 4 days after plating (100% colonies showing GFP+). This efficiency aligns with previous estimates (Wise et al., 2006). Although it is 2-3 orders of magnitude lower than electroporation (Figure 1C), it is only about 1 order of magnitude lower than that of chemically competent *Agrobacterium* cells (https://goldbio.com/product/14849/gv3101-agrobacterium-chemically-competent-cells). This level of efficiency is sufficient for routine experiments, as it allows the generation of several hundred colonies per plate using as little as 30-35 ng of DNA, and further miniaturization is feasible (Hwang et al., 2017). We used *A. tumefaciens* GV3101 strain for most tests, as it is the most efficient for *Marchantia* transformation and widely used for agroinfiltration in *N. benthamiana leaves (Li, 2011; Tsuboyama and Kodama, 2018)*. As different strains can be used for various plant transformation pipelines (Hwang et al., 2017), we tested it with three more strains: LBA4404 (Ooms et al., 1982), EHA105 (Hood et al., 1993), and AGL-1 (Lazo et al., 1991). While LBA4404 showed similar efficiency compared to GV3101, EHA105 exhibited one order of magnitude lower efficiency, and we did not obtain colonies using AGL-1. We recommend testing the efficiency before implementing high-throughput pipelines in alternative strains.

### Automation of Agrobacterium transformation

Adaptation of the freeze-thaw protocol made the process suitable for automation, and offered several advantages, including significant reductions in cost and time. In order to implement automation at low cost, we adapted the workflows for use on the *Opentrons* liquid handling robots (Figure 1B). These devices have been designed to lower the entry cost for bench-top robotic handling of liquids and maintain versatility. We initially designed four protocols using the Protocol Designer tool (https://designer.opentrons.com/): a stock preparation aliquoting protocol, two transformation protocols for 24 and 96 samples, and a plating protocol for 96 samples (Supplemental File 1-4).

Unlike previous methods for bacterial transformation using the OT2 where plating was done with droplets (Storch et al., 2020), we employed the same 6-well plate configuration used for the manual freeze-thaw method. This setting reduced the risk of cross-contamination and was useful for downstream plant transformation pipelines. In the transformation protocols, we utilized the Opentrons Gen-1 Thermocycler module (https://opentrons.com/products/modules/thermocycler/) for the heat-shock step, while manually snap-freezing in liquid nitrogen. Nonetheless, any thermocycler external to the Opentrons can be used.

Workflows were designed to handle up to 24 transformations in one run, including plating, or up to 96 with the plating step carried out as a separate protocol. Still, workflows created using the Protocol Designer tool lack flexibility and need to be adjusted to plastic labware, sample number, and volume desired. Therefore, we implemented the protocols in Jupyter Notebooks, which can be more flexibly edited and provide a useful medium to have both protocol instructions and execution commands in the same file. For this, three protocols were developed (Supplementary File 5-7) capable of executing the same functions after tailored input of the relevant parameters.

To assess the risk of cross-contamination between different plasmids during operation, we conducted a test using a 96-well plate containing alternate wells with plasmids expressing sfGFP or lacZ under constitutive bacterial promoters and observed the fluorescence. After 3 days at 28 °C, no contamination or cross-contamination was detected (Figure 1F). Additionally, the efficiency of transformations carried out using the OT2 robot exhibited similar results compared to manual manipulation (Figure 1C).

### Generation of Marchantia sporelings

The transformed bacteria obtained from the previous step can be used for further applications in *Agrobacterium*-based plant transformation protocols. For stable plant transformation, we optimized a protocol for *M. polymorpha* sporeling transformation. Sporeling transformation has been successfully applied to several accessions, such as the Tak-1/2 and Cam-1/2. Cam-1/2 is very efficient for generating spores in laboratory conditions and has been used for synthetic biology applications (Sauret-Gueto et al., 2020). Dried spores can be stored at -80 °C for years or at 4 °C for several months to years. Although spores could be produced and collected in axenic conditions, surface sterilization was used to prevent occasional fungal or bacterial contamination. Spores were sterilized by brief treatment with Milton Mini sterilization tablets (Sauret-Güeto et al. 2020)..

Previous protocols involved pre-culturing spores in either liquid media supplemented with sucrose (Ishizaki et al., 2008), solid media supplemented with sucrose (Tsuboyama and Kodama, 2014) or without sucrose (Sauret-Gueto et al., 2020). To compare both methods, we tested germination rates using various media. We found that sporelings germinated at similar rates in solid or liquid 0.5x Gamborg’s B5 without any supplement, but the germination rate was lower when supplements and sucrose were added to the liquid media (Figure 2C). Sucrose and supplements are essential during co-culture (Kubota et al., 2013; Tsuboyama and Kodama, 2014), thus it is convenient to separate the spore pre-culture media from the co-culture media as described before (Sauret-Gueto et al., 2020) to optimize germination while minimise contamination risk associated with sucrose. This is also helps preventing clumping of spores, which was often observed in sporeling liquid culture.

**Figure 2.**
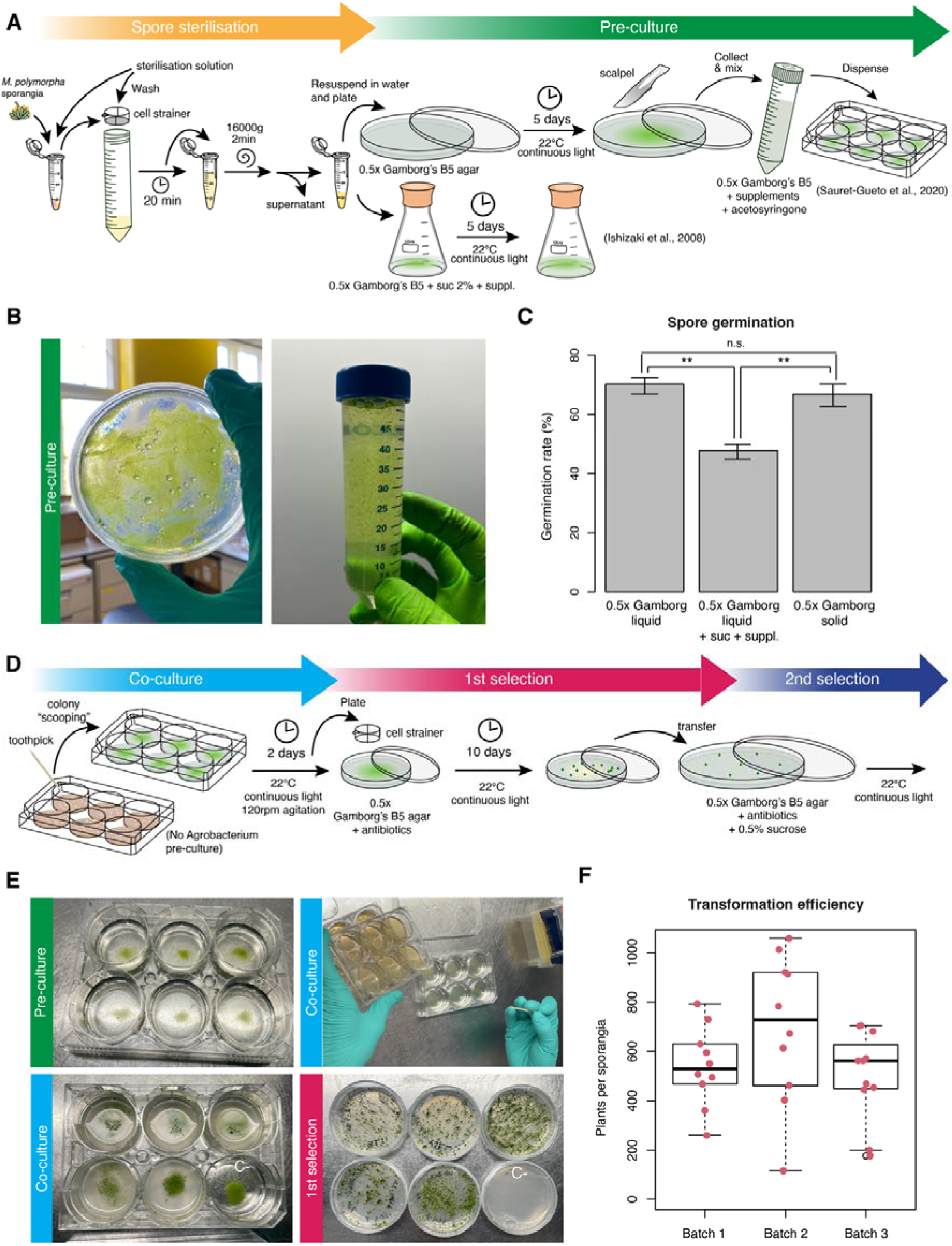
(A) Schematic diagram of Marchantia spore sterilization and pre-culture steps, including paths for germinating spores in solid media. (B) Picture of sporelings after 5 days of germination on solid medium containing 0.5x Gamborg’s B5 medium (left). Spores after collection in liquid media (centre). Spores after dispensed in a 6-well plate (right). (C) Quantification of spore germination rate in three different conditions (n > 600). Asterisks indicate statistical significance (Pairwise Binomial Exact Test; ** p-value < 0.01 n.s. p-value > 0.05) Error bars represent 95% confidence intervals. (D) Schematic diagram showing the steps for Marchantia sporeling co-culture and selection. (E) Picture of Agrobacterium colonies being inoculated into sporelings for co-culture (top, left). Picture of sporelings after 2 days of co-culture (top, right). The culture looks cloudy after co-culture, except for the lower-right well which was used as a control and not inoculated. Sporelings being collected after co-culture for selection (bottom, left). Sporelings after 10 days of selection (bottom, right). The lower-right well correspond to the same empty control used in top-right picture without inoculation. (F) Box plot of Marchantia transformation efficiency from three different batches of spores after hygromycin selection. Individual values are shown (n = 10).

### Transformation of Marchantia

Spores germinated in solid media (Figure 2B) and then collected and transferred to co-culture media (with supplements and acetosyringone). We simplified and miniaturised this step by growing spores in solid 0.5x Gamborg B5 media (figure 2A) and using 1-2 sporangiophores for up to 24 transformations (about 10 sporangia). For this purpose, we used 4 mL co-cultures media in 6-well plates as described in Sauret-Gueto et al. (2020), which provided a degree of miniaturization compared to the protocols of Ishizaki et al. (2008) (Figure 2B).

In previous protocols, the next step involved the pre-culture of transformed *Agrobacterium* strains prior to co-culture with plant material. Typically, bacteria were grown in LB liquid media for two days, followed by centrifugation, and then another growth step in 0.5x Gamborg’s B5 media with sucrose, vitamins, and acetosyringone for 6 hours before co-culture (Ishizaki et al., 2008; Sauret-Gueto et al., 2020) or for 2 days on solid media (Tsuboyama and Kodama, 2018). To simplify this part of the protocol, we tested whether *Agrobacterium* could be directly inoculated into a spore culture in liquid media. We found that “scooping” a colony from the *Agrobacterium* selection plate using a sterile toothpick, or pipette tip, straight into the co-culture worked as well as pre-culturing (see Supplemental Video 2), reducing notably hands-on work and time and the number of steps. We recommend growing the *Agrobacterium* on plates for 3 days, as colonies were easier to collect. The use of 6-well plates for *Agrobacterium* and co-culture also significantly helped with the manual handling of the experimental materials (Figure 2D).

The co-culture was then carried out for 2 days as optimized before (Ishizaki et al., 2008; Sauret-Gueto et al., 2020). After co-culturing, sporelings were collected using a 70 μm cell sterile strainer, washed, and then plated on 60 mm petri dishes. The use of small 60 mm petri dishes significantly reduces the footprint in growth chambers. The petri dishes contained selective antibiotics and cefotaxime to eliminate growth of any remaining *Agrobacteria* during the selection (Figure 2E). The sporelings were spread on the plate with a gentle circular movement (see Supplemental Video 3).

We observed an average transformation efficiency of about ∼600 transgenic plants per sporangium used among different batches with the *Marchantia Cam-1/2* strain (Figure 2G). This is comparable to the efficiency described for the Tak-1/2 strain in previous protocols under optimal conditions (Ishizaki et al., 2008; Ishizaki et al., 2015). We also observed that the Tak-1/2 strain work as well in our protocol. Overall, this demonstrates that the simplification steps made in Sauret-Gueto *et al*. (2020) and here did not have a significant negative impact on plant transformation efficiency.

For lower throughput experiments or laboratories without a supply of sporangia, a similar set-up in 6-well plates and direct inoculation of *Agrobacterium* could be used to transform thallus fragments, adapting the protocol from Kubota et al. (2013), at the likely expense of a lower efficiency of transformation.

### Optimized selection of transgenic lines

After co-cultivation and 10 days of growth, the sporelings grew large enough to be transferred to a second selection plate. This second selection step is indispensable to prevent crowding of spores, to accelerate plant growth, and to eliminate false positives. We recommend transferring 6-7 independent lines to 90 mm petri dishes to screen for genetic variability between different T-DNA insertions and avoid overcrowding (Figure 3A). Following this optimized pipeline, one could routinely transform 24 transgenic lines per week, allowing testing of up to a hundred different constructs per month with overlapping pipelines. The total footprint for two consecutive batches of transformation was less than 0.6 m^2^ (Figure 3A), significantly reducing the cost of energy and space required in growth cabinets compared to flowering plant models.

**Figure 3.**
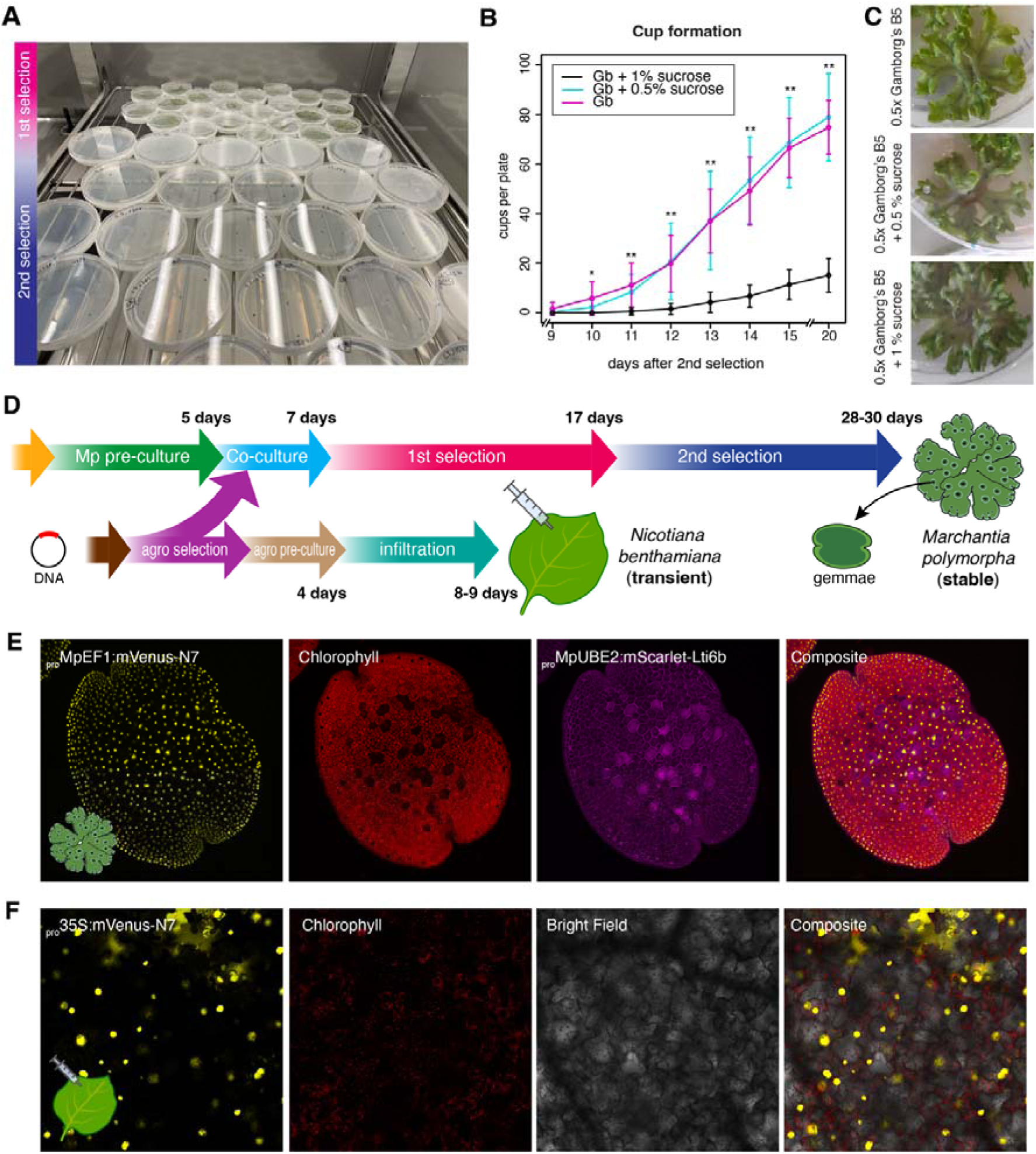
(A) Picture of two batches of Marchantia sporeling transformation showing plates growing at 1^st^ and 2^nd^ selection stage. (B) Time-course quantification of gemma cup formation in three 0.5x Gamborg’s B5 media supplemented or not with sucrose (see legend) during 1^st^ selection (n = 8). Asterisks indicate statistical significance (Pairwise T-Test between either 0.5 % (w/v) sucrose or 1% (w/v) sucrose against no sucrose at each point of the time course; * p-value < 0.05, ** p-value <0.01.). No statistical difference (p-value > 0.05) was found between 0.5% and 1% sucrose. Error bars represent SD. (C) Pictures of plants after 4 weeks in each condition tested. (D) Schematic diagram of high-throughput workflows for Agrobacterium transformation and Marchantia stable transformation and N. benthamiana leaf transient expression. (E) Example of confocal images of a Marchantia gemma transformed with a plasmid contain two fluorescent proteins constitutively expressed located in the nuclei (_pro_MpEF1a:mVenus-N7) and the plasma membrane (_pro_MpUBE2:mScarlet-Lti6b). (F) Example of a leaf of N. benthamiana transformed with a plasmid containing a fluorescent protein constitutively expressed located in the nuclei (_pro_35S:mVenus-N7). Fluorescent channels are shown separated. Scale bar = 100 μm.

The gemma of *Marchantia* is an extraordinary tissue for plant biology (Kato et al., 2020; Romani et al., 2023). Gemmae are clonal propagules that are produced in cups, and form part of a short vegetative life cycle. They can be directly used for the visualization of fluorescent reporters or other assays, providing information about the entire organism (Figure 3E). However, obtaining gemma cups can sometimes be a bottleneck for systematic pipelines, particularly in the Cam-1/2 *Marchantia* background. It has been shown that supplementation with sucrose can boost the production of gemmae cups (Terui, 1981). To address this, we tested different concentrations of sucrose to boost gemma cup production from sporelings after the first selection. We found that supplementation with 0.5% or 1% sucrose (w/v) significantly increased cup production after 10 days of selection. By day 15 of the second selection, most replicates had cups in all independent lines when supplemented with sucrose, while unmodified 0.5x Gamborg’s B5 showed no cups (Figure 3B). As there was no significant difference between 0.5% and 1% sucrose supplementation, we adopted 0.5% as the standard for our experiments due to its negligible effect on other aspects of growth (Figure 3C).

Elements of this streamlined pipeline could be implemented not only in novel model systems like *Marchantia* (Figure 3D; Figure 3E) but also in more established and rapid pipelines like agroinfiltration in *N. benthamiana* (Figure 3F). Notably, the average time to obtain stable transgenic plants using this pipeline is only three times more than doing experiments in transient systems, making both systems feasible for high-throughput experimentation.

## DISCUSSION

In this work, we have taken significant steps towards optimizing *Agrobacterium*-based transformation pipelines for high-throughput experiments *in planta*. We introduced simplified steps for freeze-thaw transformation of *Agrobacterium*, sample handling, and *Marchantia polymorpha* transformation to facilitate cost-effective and scalable high-throughput experiments without the need for specialized lab equipment. Thanks to this pipeline, a single worker can maintain an output of up to a hundred independent *Marchantia* plants transformed per month and we have successfully employed this for the screening of an arrayed library of about ∼360 transcription factor promoters (Romani et al., 2024). Further, we successfully adapted our simplified *Agrobacterium* transformation protocol to a semi-automated open source robotic platform, Opentrons OT2, aiding the development of yet higher-throughput plant transformation methods. Moreover, different versions of the protocols were created using the Protocol Designer tool and via Python code with custom Jupyter Notebooks. This makes the protocols open to scientists with different levels of programming expertise. These automated protocols have the potential to significantly reduce costs, human error, and experimental preparation time, benefitting both small laboratories and specialized biofoundries.

While automated protocols for *Escherichia coli* transformations in cloning already exist (Storch et al., 2020), there is a lack of similar methods for *Agrobacterium*. This protocol is a much-needed resource for the plant bioscience community, especially in the field of plant synthetic biology. By using the open-source Opentrons platform, we ensure that automation is accessible to a broader range of researchers without requiring additional specialised equipment. This can be coupled with high-throughput design and assembly of genetic constructs (Cai et al., 2020; Bryant et al., 2023) to make a more comprehensive pipeline. Further advances in automation and instruments could lead to fully automated protocols. We support the existing protocols with in-depth tutorials for the entire workflow in OT2 robots, reflecting our commitment to accessibility and future development.

Additionally, we have simplified and optimised *M. polymorpha* transformation and the selection of transgenic lines, utilizing this extraordinary model plant for high-throughput experiments. *Marchantia* has also been demonstrated as suitable for transient transformation using *Agrobacterium* (Iwakawa et al., 2021) or particle bombardment (Konno et al., 2018; Westermann et al., 2020), allowing for testing of subcellular localization, *in vivo* sgRNAs for CRISPR/Cas9 testing, or protein-protein interactions.

Our pipeline highlights the scalability of *Marchantia* stable transformation and the feasibility of obtaining stable isogenic transgenic lines in a cost-effective manner and within a relatively short timeframe. The bottleneck in this pipeline is no longer in generating stable transgenic lines, but rather in testing them. This demands simple read-outs, such as examining expression patterns in the gemma (Romani et al., 2023). Both *Marchantia* and *N. benthamiana* are extraordinary chassis for bioproduction and metabolic engineering and can be optimized and engineered for different applications (Golubova et al., 2024; Tansley et al., 2024; Tse et al., 2024). We envision that this pipeline will facilitate large-scale screening *in planta* and other types of screens such as functional genetics studies, enzyme activity analyses, directed evolution studies, or investigations into molecular interactions. Our optimized pipeline opens new possibilities for accelerating research in plant biotechnology and synthetic biology fields, ultimately contributing to advancements in agricultural and biotechnological applications.

## Supporting information

Supremental Files

## ACKNOWLEDGEMENTS

This work was funded by BBSRC BB/T007117/1 to J.H, BB/W014173/1 to J.H. and J.C.M., and BBSRC BB/F011458/1 for confocal microscopy. F.R. is a Leverhulme Early Career Fellow (ECF-2023-534) funded by the Leverhulme Trust and the Isaac Newton Trust (23.08(f)). We thank Adrienne Pate for providing plantlets of *N. benthamiana* and help in the infiltration experiments.

## MATERIALS AND METHODS

### Plant Growth Conditions and spore production

Spores from *Marchantia polymorpha subs. rudelaris* accessions *Cam-1* (male) and *Cam-2* (female) were utilized in this study (Delmans et al., 2017). Under normal conditions, plants were grown on solid 0.5× Gamborg’s B5 basal medium (Phytotech #G398) at pH 5.8 with 1.2% (w/v) agar of micropropagation grade (Phytotech #A296) under continuous light at 21°C with a light intensity of 100 μmol/m^2^/s.

For spore production, plants were grown in TP5000+TPD5000 Microbox^TM^ in axenic conditions on Jiffy-7 dehydrated peat disks under continuous light for a month and then moved to a station with far-red supplemented light. After a further month, sexual organs had developed and were manually fertilised using sterilised RO filtered water as transfer medium between male and female organs (Sauret-Gueto et al., 2020). Spores were collected and stored into single Eppendorf tubes, or 50mL Falcon tubes, with silica beads at 4°C.

*Nicotiana benthamiana* seeds were sown and grown for three weeks in Levington M3 compost (Scotts, Surrey, UK) within a Conviron growth chamber (Manitoba, Canada) under long-day conditions (16/8-hour light/dark cycle) at 22°C, 60% relative humidity, and 200 μmol/m /s. Agroinfiltration assays were performed on the third or fourth true leaves of the *N. benthamiana* plants.

### Plasmid used

A binary vector bearing the kanamycin resistance gene and containing the bacterial promoter pJ23106 driving expression of sfGFP (pCk1-ye) or lacZ (pCk1) was used (Sauret-Gueto et al., 2020). For *Marchantia* and *Nicotiana benthamiana* transformation, the plasmids *_pro_MpEF1a:mVenus-N7; _pro_MpUBE2:mScarlet-Lti6b* (pBy01 backbone) and *_pro_Mp35S:mVenus-N7; _pro_MpUBE2:mScarlet-Lti6b* (pBy01 backbone) were used respectively (Romani et al., 2023). The _pro_MpUBE2 showed no activity in *N. benthamiana*.

### Agrobacterium tumefaciens transformation

*A. tumefaciens* strain GV3101 was performed as described before (Weigel and Glazebrook, 2006) for electroporation and freeze-thaw transformation for strains GV3101, EHA105, AGL-1 and LBA4404 as described below in detail (Supplementary Document 1). Images were captured using a GelDoc (BioRad). Transformation efficiency was calculated by estimating the amount of DNA used with a NanoDrop One microvolume UV-VIS spectrophotometer (Thermo Scientific) and counting CFU 3 days after plating for at least 3 independent transformations.

For each protocol, we used Gen-2 p300 and p20 pipettes Opentrons pipettes, along with Opentrons pipette tips for their respective pipettes. We used skirted 96-well plates (BioRad #HSP9601) and 6-well plate flat-bottom cell culture plates (Greiner #657160) unless something else is specified. The protocols also use the Opentrons gen-1 thermocycler and gen-2 temperature modules.

For the automated workflow using the Opentrons OT-2 platform and Protocol Designer we adapted the same steps described for the manual protocol. The *Agrobacterium* strain was grown as described in detail in Supplementary Document 1. We created a protocol for the aliquoting of the competent cells in 96-well plates (Supplementary File 1), and subsequent files for plasmid transformation using up to 24 1.5 mL Eppendorf tubes for the plasmid DNA Supplementary File 2) or a 96-well source plate (Supplementary File 3). After the incubation, cell plating was implemented as a separate protocol that can handle up to 96 transformations (Supplementary File 4). The layout and outline of the automated pipeline are shown in Supplemental Figure 1.

Each of the files has instructions to be followed along with the execution of the protocol, but general instructions on how to run, edit and simulate OT2 protocols can be found on our dedicated automation protocols website (https://openplant.github.io/openplant_automation_protocols). The topics addressed by the tutorials span basic execution of protocols with the Opentrons app through to advanced scripting for protocols with the Python API. To run the Jupyter Notebook protocols, refer to Tutorials > <u>Using Jupyter Notebooks with the OT2</u>.

### Spore germination and gemma cup counting

Spores were cultured in normal growing conditions on solid plates or in liquid culture containing only 0.5x Gamborg’s B5 without agar or supplemented with 2% (w/v) sucrose (Fisher), 0.03% (w/v) L-glutamine (Alpha-Caesar), and 0.1% (w/v) casamino acids (Fisher). Liquid cultures were grown under the same light and temperature conditions with 120 rpm agitation. Spore germination was assessed as the ratio between chlorophyll-containing spores and dead spores after 6 days using a fluorescent stereomicroscope Leica M205 FA. For quantifying gemma cup formation, plants were grown for 10 days in normal selection conditions and then 6 plants were transferred to a plate with 0%, 0.5%, 1% (w/v) sucrose for 20 days and cups were counted every day after the first cup appeared (9 days after second selection) in 3 biological replicates.

### Agroinfiltration of *N. benthamiana* leaves

Agroinfiltration was performed as previously described (Li, 2011). Briefly, 6 ml of LB and antibiotics were inoculated with *Agrobacterium tumefaciens* GV3101 grown overnight at 28°C shaking and 220 rpm, with the concentration normalized to O.D. 1.4. Bacterial cultures were centrifuged at 4500 g for 10 min, resuspended in resuspension media (10 mM MgCl2, 10 mM MES pH 5.6), and infiltrated using 1 mL syringes on the ventral surface of *N. benthamiana* leaves. Visualization was performed 4 days post-infiltration.

### *Marchantia* transformation and selection

*Agrobacterium*-mediated transformation of M. polymorpha was carried out following a modified version of the previous 6-well plate method (Sauret-Gueto et al., 2020; Romani et al., 2024). Briefly, *Marchantia* sporelings were sterilized and cultures in solid 0.5x Gamborg’s B5 plates for 5 days and collected in 6-well plates (Greiner #657160) of liquid 0.5x Gamborg’s B5 with supplements. Colonies of *Agrobacterium* were picked with a sterile tip or toothpick, incubated into the wells, and growth for 2 days in agitation. Spores were collected in solid 0.5x Gamborg’s petri dishes with antibiotics and selected for 10 days. A second round of selection was carried out in solid 0.5x Gamborg’s with antibiotics supplemented or not with sucrose. Details of the protocols are explained and illustrated in Supplementary Document 2.

### Confocal microscopy

Confocal images of *Marchantia* and *N. benthamiana* were acquired using a Leica SP8X spectral confocal microscope upright system equipped with a 460−670 nm super continuum white light laser, 2 CW laser lines at 405 nm and 442 nm, and a 5 Channel Spectral Scanhead (4 hybrid detectors and 1 PMT). Imaging was conducted using a 10× air objective (HC PL APO 10×/0.40 CS2). The excitation laser wavelength and the captured emitted fluorescence wavelength window were as follows: for mVenus (514 nm, 527−552 nm), for mScarlet (561 nm, 595−620 nm), and for chlorophyll autofluorescence (633 nm, 687−739 nm).

---

**Supplemental Figure 1.**
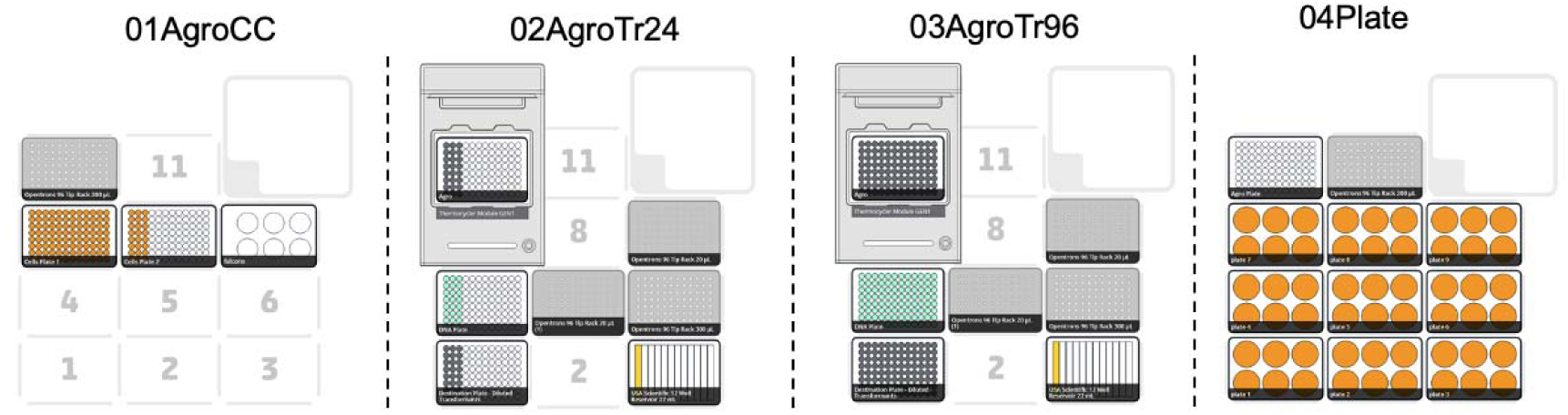
Layout of different protocols for automated *Agrobacterium* transformation protocols.

**Supplemental Video 1.** Video of manual *Agrobacterium* transformation plating

**Supplemental Video 2.** Video of *Marchantia* co-culture.

**Supplemental Video 3.** Video of *Marchantia* plating.

**Supplemental File 1.** Protocol Designer file for *Agrobacterium* competent cells dispensation in 96-well plates.

**Supplemental File 2.** Protocol Designer file for *Agrobacterium* transformation automation for 24 samples.

**Supplemental File 3.** Protocol Designer file for *Agrobacterium* transformation automation for 96 samples transformation.

**Supplemental File 4.** Protocol Designer file for plating *Agrobacterium* transformations.

**Supplemental File 5.** Jupyter Notebook protocol for *Agrobacterium* transformation.

**Supplemental File 6.** Jupyter Notebook protocol for plating *Agrobacterium* transformations.

