## Supplementary material for "Semi-automated workflow for high-throughput *Agrobacterium*-mediated plant transformation": Supremental Files: 01Agrobacterium Transformation.ipynb

````
# %% [markdown]
## Protocol Initialization
Ensure you have calibrated the pipettes before doing this protocol

## Before you proceed
This protocol requires the use of the thermocycler lid, which is not easily accessible through jupyter notebooks.

To be able to access the lid, go to the notebook file system tab, then go to New > Terminal and run the command:

```
systemctl stop opentrons-robot-server
```

This will disconnect the robot from the opentrons app so that only this protocol is able to access it. This is also explained in our [Jupyter Notebook protocols tutorial](https://openplant.github.io/openplant_automation_protocols/Tutorials/Jupyter%20Notebooks/)

To re-connect the robot to the app, run:

```
systemctl start opentrons-robot-server
```

##Furthermore remember that the file "azenta_96_wellplate_200ul.json" and "enzymax_12_reservoir_20ml.json" must be storred in the same directory for this code to work


# %% [markdown]
#convert this cell to "code" if you haven't installed the opentrons library yet, you might want to use !conda instead of !pip to install this package if you work in a conda environment
!pip install --upgrade opentrons 

#chose if you are executing the protocol with a robot or simulating it o your computer by choosing one of the next two cells accordingly


# %% [code]
#if executing on a robot:
import opentrons.execute
from opentrons import protocol_api
import json

# %% [code]
protocol = opentrons.execute.get_protocol_api('2.13')
protocol.home()
protocol.set_rail_lights(True)

# %% [markdown]
#if simulating:
from opentrons import simulate
import json

protocol = opentrons.simulate.get_protocol_api('2.13')
protocol.home()

# %% [markdown]
it might give you a warning like "/Users/user_name/.opentrons/robot_settings.json not found. Loading defaults" but don't worry about it and keep going

# %% [markdown]

## Declaring Labware
(in reverse order)
- **Left mount**: An 8-channel p20 pipette
- **Right mount**: An 8-channel p300 pipette
- Thermocycler module plus 96-well PCR plate on it
- **Slot 3**: 12-well reservoir for LB
- **Slot 1**: Destination plate (96-well PCR plate)
- **Slot 4**: DNA plate (96-well PCR plate)
- **Slots 6 and 9**: Tipracks for both pipettes

Replace with appropriate labware codes in the cell below if needed
If you need to use custom labware, use our [Jupyter Notebook protocols tutorial](https://openplant.github.io/openplant_automation_protocols/Tutorials/Jupyter%20Notebooks/)

# %% [code]
tipracks_20 = [protocol.load_labware('opentrons_96_tiprack_20ul', '5'), protocol.load_labware('opentrons_96_tiprack_20ul', '9')]
tipracks_300 = [protocol.load_labware('opentrons_96_tiprack_300ul', '6')]

dna_plate = protocol.load_labware('biorad_96_wellplate_200ul_pcr', '4')
destination_plate = protocol.load_labware('biorad_96_wellplate_200ul_pcr', '1')

# Here using 12-well reservoir for a ma of 20mL.
# Since a 96-well run would use almost 18mL, it might be advisable to use a 1-well reservoir 
with open('enzymax_12_reservoir_20ml.json') as labware_file:
    labware_def = json.load(labware_file)
    lb_res = protocol.load_labware_from_definition(labware_def, 3)
    
thermocycler = protocol.load_module('thermocycler module')

p20_8 = protocol.load_instrument('p20_multi_gen2', 'left', tip_racks=tipracks_20)
p300_8 = protocol.load_instrument('p300_multi_gen2', 'right', tip_racks=tipracks_300)

# %% [markdown]
# Define the number of samples here
- Run the next cell and fill out the prompts
- You can use from 1 to 96 samples. Do note that using non-multiples of 8 will still make the robot use a whole column of pipette tips.

# %% [code]
samples_no = int(input("Number of samples: "))
final_volume = int(input("What volume do you want the cells to be in at the end? (in ul, max 200) "))
do_dilution = str(input("Do you want to dilute the cells at the end of the protocol? (Y/N) "))
if do_dilution in ["Y", "y"]:
    dilution_amount = int(input("How much of the cells do you want to pick to dilute? (in ul, max 50) "))
else:
    print('You chose not to dilute the cells at the end of the protocol')

if samples_no - 1 in range(96):
    columns_no = samples_no//8
    if samples_no % 8 > 0:
        columns_no += 1
else:
    samples_no = 0
    print('Invalid number of samples')

# %% [code]
tubes = int(input("Do you want to use PCR strips or 96 well plates? Answer '1' for strips or '2' for plates: "))
print(tubes)
#the biorad 96 wel plate from the opentron library perfectly fits into the pcr mtal plate, but if you use PCR 
# they are usually taller than tht and the OT2 needs to know it or will not go high enough when moving form one
#to the other, and will crash everything.

if tubes==1:
    with open('azenta_96_wellplate_200ul.json') as labware_file:
        PCR_strip_def = json.load(labware_file)
        cells_plate = thermocycler.load_labware_from_definition(PCR_strip_def)
        print("loading on PCR succesful")
else:
    cells_plate = thermocycler.load_labware('biorad_96_wellplate_200ul_pcr')

# %% [markdown]
## Labware Calibration

Change the x, y, and z variables (**in mm**) in the "set_offset" functions and run each cell repeatedly until the positioning is correct for that piece of labware. You will have to use a "dummy" plate to calibrate the labware that is not on the deck yet (competent cells).

### Tipracks:

# %% [code]
protocol.comment('--- CALIBRATION ---')

# %% [code]
tipracks_20[0].set_offset(x=-0.3,y=0.8,z=0.0)
p20_8.move_to(tipracks_20[0]['A1'].top())

# %% [code]
tipracks_20[1].set_offset(x=0.2,y=0.8,z=0.0)
p20_8.move_to(tipracks_20[1]['A1'].top())

# %% [code]
tipracks_300[0].set_offset(x=0.8,y=0.7,z=0.0)
p300_8.move_to(tipracks_300[0]['A1'].top())

# %% [markdown]
### Plates/Racks:

# %% [code]
p20_8.pick_up_tip()

# %% [code]
dna_plate.set_offset(x=0.0,y=0.2,z=0.0)
p20_8.move_to(dna_plate['A1'].top())

# %% [code]
destination_plate.set_offset(x=0.0,y=0.3,z=0.5)
p20_8.move_to(destination_plate['A1'].top())

# %% [code]
cells_plate.set_offset(x=0.0,y=0.3,z=0.0)
p20_8.move_to(cells_plate['A1'].top())

# %% [code]
lb_res.set_offset(x=0.0,y=0.0,z=0.0)
p20_8.move_to(lb_res['A1'].top())

# %% [code]
p20_8.return_tip()
p20_8.reset_tipracks()
protocol.home()

# %% [markdown]
## Protocol Execution
Flash freeze and heat shock

# %% [code]
print("Setting temperature to 4°C")
thermocycler.set_block_temperature(temperature=4)
print('Done')

# %% [markdown]
- Place the frozen competent cells on the thermocycler block.
- Run the next cell immediately

# %% [code]
protocol.comment('--- TRANSFERRING DNA INTO COMPETENT CELLS ---')
print(protocol.commands()[-1])
print('Waiting 10 minutes...')
protocol.delay(minutes=10)
for col in range(columns_no):
    print('Transferring column', col + 1)
    p20_8.pick_up_tip()
    p20_8.aspirate(5, dna_plate.rows()[0][col])
    p20_8.dispense(5, cells_plate.rows()[0][col].bottom(z=1.5))
    p20_8.mix(3, 5, cells_plate.rows()[0][col])
    p20_8.touch_tip()
    p20_8.drop_tip()
print("Done")

# %% [markdown]
- Flash freeze the plate with cells
    - **REMEMBER** to flick the tubes before freezing
- Run the next cell immediately, while you freeze the cells

# %% [code]
protocol.comment('--- PREPARING TEMPERATURE FOR HEAT SHOCK ---')
print(protocol.commands()[-1])
print('Setting block and lid temperature to 37°C')
thermocycler.set_block_temperature(temperature=37)
thermocycler.set_lid_temperature(temperature=37)
print("Done")

# %% [markdown]
- Reinsert frozen cells into the thermocycler module

# %% [code]
thermocycler.close_lid()
print("Incubating at 37°C for 5 minutes")
thermocycler.set_block_temperature(temperature=37)
protocol.delay(minutes = 5)
print("Incubating at 28°C for 60 minutes")
thermocycler.set_block_temperature(temperature=28)
protocol.delay(minutes = 60)
thermocycler.open_lid()
thermocycler.deactivate()
print("Done")

# %% [code]
if do_dilution in ['Y', 'y']:
    protocol.comment('--- TRANSFERRING LB INTO DILUTION PLATE ---')
    print(protocol.commands()[-1])
    p300_8.pick_up_tip()
    for col in range(columns_no):
        print('Transferring column', col + 1)
        p300_8.aspirate(final_volume - dilution_amount, lb_res.rows()[0][0])
        p300_8.dispense(final_volume - dilution_amount, destination_plate.rows()[0][col].top()) # I removed the touch tip here, and am dispensing from the top of the well to avoid contamination
        p300_8.move_to(lb_res.rows()[0][0].top())
        p300_8.blow_out()
    p300_8.drop_tip()

    protocol.comment('--- TRANSFERRING TRANSFORMED CELLS INTO NEW PLATE ---')
    print(protocol.commands()[-1])
    for col in range(columns_no):
        print('Transferring column', col + 1)
        p300_8.pick_up_tip()
        p300_8.aspirate(dilution_amount, cells_plate.rows()[0][col])
        p300_8.dispense(dilution_amount, destination_plate.rows()[0][col].bottom(z=4.5))
        p300_8.touch_tip()
        p300_8.drop_tip()
else:
    protocol.comment('--- TRANSFERRING LB INTO CELLS PLATE ---')
    print(protocol.commands()[-1])
    p300_8.pick_up_tip()
    for col in range(columns_no):
        print('Transferring column', col + 1)
        p300_8.aspirate(final_volume - 50, lb_res.rows()[0][0])
        p300_8.dispense(final_volume - 50, cells_plate.rows()[0][col].top()) # I removed the touch tip here, and am dispensing from the top of the well to avoid contamination
    p300_8.drop_tip()
print('Done')

# %% [code]
protocol.home()
protocol.set_rail_lights(False)

# %% [code]
protocol.commands()
````
