## Supplementary material for "Semi-automated workflow for high-throughput *Agrobacterium*-mediated plant transformation": Supremental Files: 02Plating transformed cells.ipynb

```
# %% [markdown]
## Protocol Initialization
Ensure you have calibrated the pipette before running this protocol

# %% [markdown]
#convert this cell to "code" and run it if you haven't installed the opentron library yet, you might want to use "!conda" instead of "!pip" to install this package if you work in a conda environment !pip install --upgrade opentrons

# %% [markdown]
- When running the protocol with a robot, replace "simulate" below with "execute"

# %% [code]
from opentrons import simulate, execute
from opentrons import protocol_api
import json

# %% [code]
#if simulating
protocol = simulate.get_protocol_api('2.12')
protocol.home()

# %% [markdown]
# if running on the robot
protocol = opentrons.execute.get_protocol_api('2.13')
protocol.home()

# %% [markdown]
## Declaring Labware
(in reverse order)
- **Right mount**: A p300 pipette
- **Slot 10**: A 96-well PCR plate with transformed cells
- **Slot 1 to 9**: 6-well plates up to the necessary number
- **Slots 11**: Tiprack for the pipette

Replace with appropriate labware codes in the cell below if needed
If you need to use custom labware, use our [Jupyter Notebook protocols tutorial](https://openplant.github.io/openplant_automation_protocols/Tutorials/Jupyter%20Notebooks/)

# %% [code]
tipracks_300 = [protocol.load_labware('opentrons_96_tiprack_300ul', '11')]

media_plates = [protocol.load_labware('corning_6_wellplate_16.8ml_flat', str(num + 1)) for num in range(9)]

p300 = protocol.load_instrument('p300_single_gen2', 'right', tip_racks=tipracks_300)

# %% [markdown]
# Define the number of samples here
- Run the next cell and fill out the prompts
- You can use from 1 to 96 samples

# %% [code]
samples_no = int(input("Number of samples:"))
dispense_volume = int(input("What volume to dispense in each plate well? (in uL, max 200)"))
print()
changes = 0

if samples_no - 1 in range(96):
    print(samples_no, 'samples will require', (samples_no//6)+1, 'plates')
    if samples_no > 54:
        print('you will need to change the plate at one point')
else:
    samples_no = 0
    print('Invalid number of samples')

# %% [code]
tubes = int(input("Do you want to use PCR strips or 96 well plates? Answer '1' for strips or '2' for plates: "))
print(tubes)
#the biorad 96 wel plate from the opentron library perfectly fits into the pcr mtal plate, but if you use PCR 
# they are usually taller than tht and the OT2 needs to know it or will not go high enough when moving form one
#to the other, and will crash everything.

if tubes==1:
    cells_plate = protocol.load_labware('opentrons_96_aluminumblock_generic_pcr_strip_200ul', 10)
else:
    cells_plate = protocol.load_labware('biorad_96_wellplate_200ul_pcr', 10)

# %% [markdown]
## Labware Calibration

Change the x, y, and z variables in the "set_offset" functions and run each cell repeatedly until the positioning is correct for that piece of labware. You will have to use a "dummy" plate to calibrate the labware that is not on the deck yet (competent cells).

### Tipracks:

# %% [code]
protocol.comment('--- CALIBRATION ---')

# %% [code]
tipracks_300[0].set_offset(x=0.0,y=0.0,z=0.0)
p300.move_to(tipracks_300[0]['A1'].top())

# %% [markdown]
### Plates/Racks:

no calibration for the 6-well plates

# %% [code]
p300.pick_up_tip()

# %% [code]
cells_plate.set_offset(x=0.0,y=0.0,z=0.0)
p300.move_to(cells_plate['A1'].top())

# %% [code]
p300.return_tip()
p300.reset_tipracks()

# %% [markdown]
## Protocol Execution
run only the cell below repeatedly, up to the necessary number of plate changes

# %% [code]
protocol.comment('--- PLATING CELLS PASS N° '+ str(changes + 1) +' ---')
for sample in range(samples_no):
    protocol.comment('')
    cur_plate = sample//6
    p300.pick_up_tip()
    p300.aspirate(dispense_volume, cells_plate.wells()[sample + 54 * changes])
    p300.dispense(dispense_volume, media_plates[cur_plate].wells()[sample % 6].top())
    protocol.delay(seconds=3)
    p300.touch_tip()
    p300.drop_tip()
    if sample == 53:
        samples_no -= 54
        changes += 1
        print('Change plates then run this cell again')
        break

# %% [code]
protocol.home()

# %% [code]
protocol.commands()

# %% [code]
```
