## Supplemental Document 1 for "Semi-automated workflow for high-throughput *Agrobacterium*-mediated plant transformation"

**Supplemental Document 1. Detailed protocol for *Agrobacterium* freeze-thaw method**

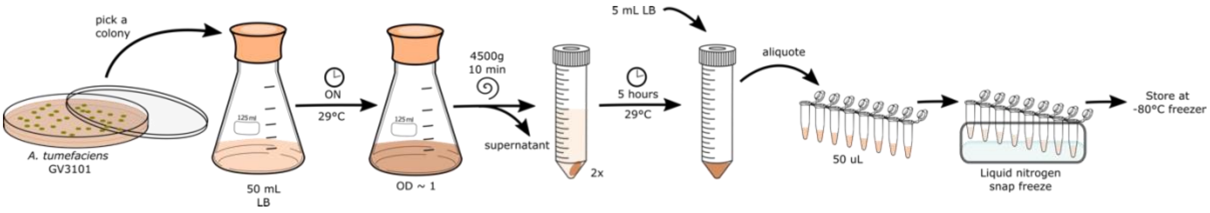


***Figure 1.*** *Diagram of Agrobacterium freeze-thaw competent cell preparation steps.*

1) Prepare competent cells. Inoculate two 50 mL Falcon tubes with 10 mL of LB with *Agrobacterium tumefaciens* strain GV3101 and grow overnight at 28 °C with shaking.
2) Pellet the cells at 4500 g for 10 min at room temperature.
3) Discard supernatant and resuspend in 1 mL LB each.
4) Aliquot 50 μL of cells in sterile 200 μL PCR tubes for direct use, or 200μL in 1,5 mL Eppendorf tubes.
5) Snap freeze in liquid nitrogen and store at -80 °C for future transformations.

*Note: Agrobacterium competent cells can be used right away without storing at -80 °C.*

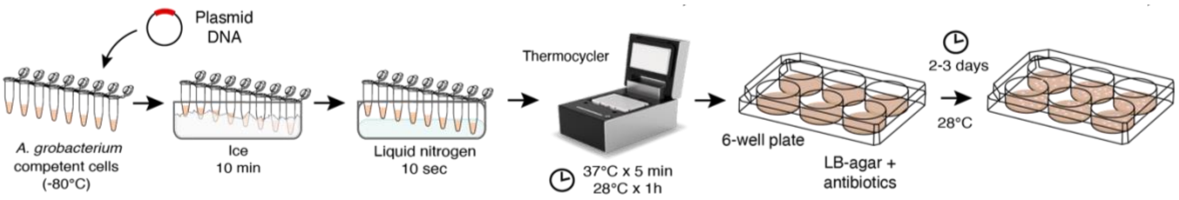


***Figure 2.*** *Diagram of Agrobacterium freeze-thaw transformation steps.*

6) Thaw frozen competent cells for 10 min in ice.
7) Use 50 μL of competent cells per transformation (may need aliquoting if stocks are stored in 1,5 mL Eppendorf tubes) in 200μL PCR tubes or a 96 well plate.
8) Add 2 μL of standard miniprep binary plasmid (100 ng/ul). Mix by gently flicking.

*Note: We recommend using one negative control with water in every batch.*

9) Flash-freeze in liquid nitrogen until it stops boiling (~10-15 sec),

10) Insert the tubes into a Thermocycler with program: 37 °C for 5 minutes, 28 °C for 1 hour.

*Note: water bath at 37 °C and incubator at 28 °C can alternatively be used.*

11) Add 50 µL of LB and plate on 6-well plates with LB with selective antibiotics for the plasmid and for the strain (for antibiotics working concentration see below). Gently spread the liquid on the plate doing orbital movements (Suppl. Video 1).
12) Culture plates upside-down at 28 °C for 3 days.

**Antibiotic working concentrations**

For a 500 ml Media bottle add:

*for bacteria*

- 500 µL of a 50 mg/mL kanamycin (Melford, #K22000) stock in water (50 µg/mL final)
- 250 µL of a 50 mg/mL rifampicin (Melford, #R64000) stock in DMSO (25 µg/mL final)
- 500 µL of a 25 mg/mL Chloramphenicol stock in ethanol (25 µg/mL final) (Sigma-Aldrich, #C0378)
- 500 µL of a 25 mg/mL gentamicyn (Duchefa, #1405-41-0) stock in water (25 µg/mL final)
- 250 µL of a 100 mg/mL spectinomycin (Melford, #S23000) stock in water (50 µg/mL final)
