## Supplemental Document 2 for "Semi-automated workflow for high-throughput *Agrobacterium*-mediated plant transformation"

**Supplemental Document 2. Detailed protocol for *Marchantia* transformation**

**Spore sterilization and pre-culture**

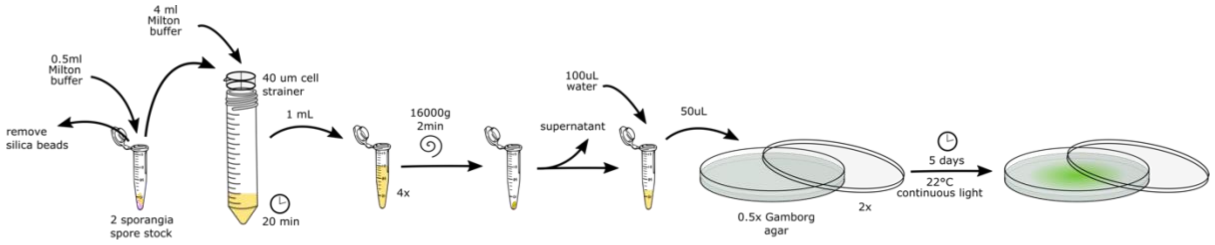


***Figure 1.*** *Diagram of spore sterilization steps.*

1) Start from 1-2 archegoniophores dried in an Eppendorf tube with silica beads.
2) Remove the silica bead from the tube or move the archegoniophore to a new tube.
3) Prepare the sterilisation solution by adding 1 Milton mini sterilizing tablets (https://www.milton-tm.com/en/consumer/products/mini-sterilising-tablets) tablet to 10 ml of sterile water in a 50 mL Falcon tube.
4) Add 200 μL of the sterilization solution in a tube with sporangia and crush and mix thoroughly with the tip to release the spores from the sporangia until the solution turns dark yellow.
5) Add 300 μL of sterilization solution and mix till homogeneous.
6) Place a 40 μM cell strainer (Greiner #542040) on a 50 mL falcon tube and pour the 0.5 mL of the yellow liquid onto the filter avoiding green tissue to avoid clogging.
7) Wash the filter with 2.5 mL of sterilization solution.
8) Divide the filtrate solution containing spores (total volume 3 mL) into two 1.5 mL Eppendorf tubes. Let samples stand for 15-40 min.
9) Prepare two 0.5x Gamborg's B5 agar plates.
10) Spin tubes at 16000 g for 2 min in a tabletop microcentrifuge.
11) Discard supernatant without disturbing the yellow pellet and pool spores in a single tube.
12) Re-suspend in ~100 μL sterile water and inoculate 50 μL on the 0.5x Gamborg's B5 agar plates agar plates and plate the spores using a sterile spreader.

*Note: that quantity works well for ~24 transformations, adjust depending on the number of transformations needed.*

13) Seal the plates with micropore tape and culture for 5 days upside-down at 22 °C in continuous light.

#### ***Agrobacterium*** **and** ***Marchantia*** **co-culture**

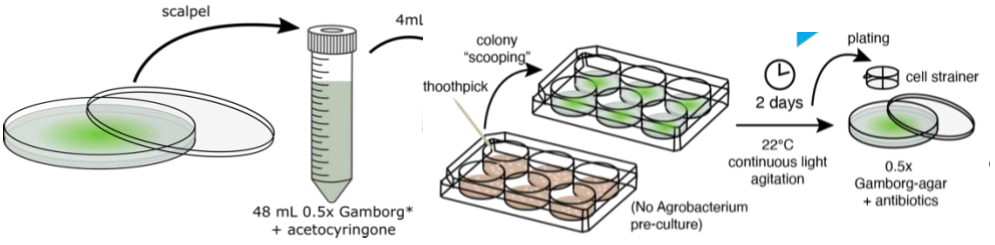


***Figure 2.*** *Diagram of co-culture steps for Marchantia transformation.*

1) Collect the spores from the agar plates with a sterile scalpel and pool them in two 50 mL Falcon tubes containing 50 mL of liquid 0.5x Gamborg's B5 media (plus supplements) and acetosyringone (Phytotech #A104) to a final concentration of 100 µM.

*Note: adjust the volume of liquid media according to the number of planned transformations.*

2) Add 4 mL to each well in four 6-well plates.
3) With a sterile tip or toothpick, scoop one colony of *Agrobacterium* with the appropriate plasmid and inoculate one well of spores (Suppl. Video 2).

*Note: make sure that most of the bacteria does not remain in the tube or tip after inoculating. We recommend leaving one well without inoculation as a control for contamination. More than one Agrobacterium strain containing plasmids with different antibiotic resistance can be co-transformed. A 1-2 orders of magnitude reduction in efficiency should be expected. For that purpose, hygromycin, G418, and chlorosulfuron are the most effective antibiotics for plant selection (Tsuboyama et al., 2018).*

4) Repeat for each well.
5) Seal the 6-well plate with double micropore tape and grow for 2 days at 22 °C with shaking and continuous light.

*Note:* co-culture should look cloudy but spores still green. If the co-culture is still clear during the first day, repeat the *Agrobacterium* inoculation.

6) Using a pipette, transfer the whole content of sporelings from each well to a 70 μm cell strainer (Greiner, #542070) in a 50 mL Falcon tube.
7) Wash with 10 mL sterile water.
8) Using 400 μL of sterile water, wash the sporelings onto 60 mm Petri dish with 0.5x Gamborg's B5 plus 100 μg/ml cefotaxime and selective antibiotic/s for the plasmid of interest (normally Hygromycin).
9) Repeat for each well.
10) When the plate is dry, add double micropore tape and culture at 22 °C with continuous light.

*Note: after 5-7 days the selection should be evident and markers visible.*

### Selection of *Marchantia* transgenic lines

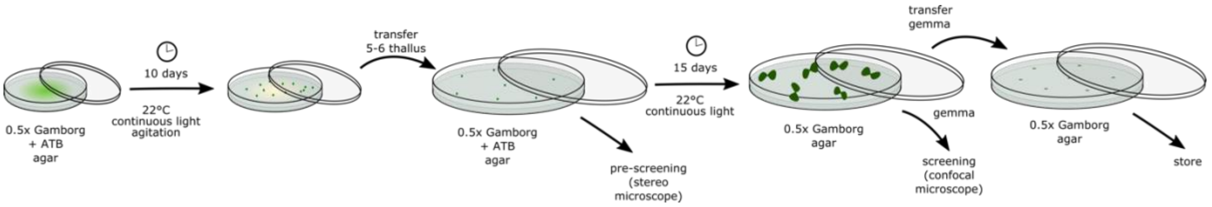


***Figure 3.*** *Diagram of selection of stable transgenic lines of Marchantia.*

11) After 10 days, using sterile tweezers, transfer 5-7 thallus to a 90mm Petri dish with 0.5x Gamborg B5 agar with cefotaxime, selecting antibiotics, and 0.5% (w/v) sucrose for a second selection in continuous light at 22 °C with.
12) When plants generate gemma cups (10-15 days later), transfer one gemma for every 4-5 lines to a final plate without antibiotics. These gemma can also be imaged for fluorescent reporters.

### Media recipes

**1L Gamborg's B5 media for sporeling transformation (liquid co-culture media)**

- 1.6 g Gamborg's B5 media (0.5x) (Phytotech, #G398)
- 20 g sucrose (2% w/v) (Fisher-Scientific S2-50GM)
- 0.30 g L-glutamine (0.03% w/v) (Sigma-Aldrich #G8540)
- 1g N-Z amine A (0.1% w/v) (Milipore #C0626)
- Fill up to 1L with RO water
- Adjust pH to 5.7-5.8 with 1 M KOH

**1L Gamborg's B5 media for *Marchantia* plates (solid)**

- 1.6 g Gamborg's B5 media (0.5x) (Phytotech, #G398)
- Mix and fill up to 1 L with RO water.
- Adjust pH to pH 5.7-5.8 with 1M KOH.
- 12 g Micropropagation-grade agar (Phytotech #A296)

Note: for 2$^{nd}$ selection add 5 g sucrose (0.5% w/v)

**Antibiotic working concentrations**

*for plants*

- 500 µL of a 100 mg/mL cefotaxime (Sigma-Aldrich, #64485-93-4) stock in water (100 µg/mL final)
- 200 µL of 50 mg/mL Hygromicin B (ThermoFisher, #10687010) (20 µg/mL final)
- 50 µL of a 500 mM chlorosulfuron (Supelco, #34322) stock in DMSO (0.5µM final)
- 50 µL of 50 mg/L G418 Sulfate (Gibco, #10131035).
